# Labellable Phylogenetic Networks

**DOI:** 10.1101/2023.02.09.527917

**Authors:** Andrew Francis, Mike Steel

## Abstract

Phylogenetic networks are mathematical representations of evolutionary history that are able to capture both tree-like evolutionary processes (speciations), and non-tree-like “reticulate” processes such as hybridization or horizontal gene transfer. The additional complexity that comes with this capacity, however, makes networks harder to infer from data, and more complicated to work with as mathematical objects.

In this paper we define a new, large class of phylogenetic networks, that we call *labellable*, and show that they are in bijection with the set of “expanding covers” of finite sets. This correspondence is a generalisation of the encoding of phylogenetic forests by partitions of finite sets. Labellable networks can be characterised by a simple combinatorial condition, and we describe the relationship between this large class and other commonly studied classes. Furthermore, we show that all phylogenetic networks have a quotient network that is labellable.

## 1. Introduction

The problem of describing the way that a set of organisms are related through evolution is usually answered by presenting a phylogenetic tree or network whose leaves are labelled by the set of species. These are directed acyclic graphs with a single root that is a common ancestor of the set of species. The internal vertices represent historical events that include speciaton, in which a vertex has out-degree two or greater, or some form of reticulation (hybridization or horzontal gene transfer), in which a vertex has in-degree two or greater. Phylogenetic trees are the special case of networks for which all internal vertices have in-degree 1, and so the only event represented by the vertices is speciation.

Manipulating either trees or networks mathematically is important because many methods for determining a tree or network require searching to find an optimum, and often they involve choosing random trees or networks. To make this possible, encodings of such graphs in other mathematical formats is often important computationally. For instance, a classical encoding of trees is the Newick format, which records clusters (descendents of internal vertices) in a structured, in-line notation (used in [7]), and there are encodings using sequences of integers [1, 13]. In networks, encodings may require additional structure, such as the use of ‘circular’ permutations for a class of *unrooted* phylogenetic networks [8].

In this paper we will show that a large class of rooted phylogenetic networks is encoded by certain covers of finite sets. This specifically generalises encodings for phylogenetic trees and forests that were given for binary trees by Diaconis and Holmes in 1998 [4], and for general phylogenetic forests more recently [10].

This encodable class we call the *labellable* phylogenetic networks, because they have the property that their internal vertices can be deterministically labelled via an algorithm based on an approach developed for trees [6].

There are many classes of phylogenetic network, and new ones are regularly defined. The reason is that inference of networks is a difficult problem, partly because the space of all networks is too large, and many networks have features that are unlikely to be inferable from data, or that make them difficult to work with mathematically. The class of labellable networks is a new and large class, as we will show in Section 6. Its most important property — that its elements are in bijection with the set of *expanding covers* (we define this later) — means that these networks can be studied using an elementary combinatorial (set-theoretic) structure. Surprisingly, we are able to show that every phylogenetic network has a labellable quotient.

The paper begins with a background section giving formal definitions and recalling some relevant results from earlier work. Section 3 describes an algorithm for labelling internal vertices of a network, and defines a labellable network as one for which that algorithm is well-defined. It also provides a structural characterisation of labellable networks, in Theorem 3.3. The next section (Section 4) gives the connection to covers, and proves that the set of labellable networks is in bijection with the set of ‘expanding covers’ (Theorem 4.4). This correspondence is made explicit for non-degenerate networks in Section 5. We then consider in Section 6 the class of labellable networks and its relationships with other well-known classes, such as tree-based networks, tree-child networks, orchard networks, and others. We show that the class of labellable networks contains all orchard networks, and hence all tree-child and normal networks, but it does not contain, and is not contained in, the class of tree-based networks. We characterise exactly when a binary tree-based is also labellable, in Theorem 6.3. Fig. 6 shows the relationship among some of these classes. Finally, in Section 7, we show that by defining an equivalence relation on vertices in any phylogenetic network, we can form a quotient network that is labellable. We call the quotient the *derived* network, and briefly discuss the relationship of this quotient to the normalisation map that was recently defined [9]. We end with a discussion that highlights some questions for further research.

## 2. Background

### 2.1. Phylogenetic trees and networks

A *(rooted) phylogenetic network* on *n* leaves is a directed acyclic graph (DAG) that has a single root (vertex with indegree 0) and *n* leaves (vertices of out-degree 0 and in-degree 1). Vertices that are not leaves or the root are called *internal*. The set of rooted phylogenetic networks on *n* leaves is denoted 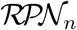. In this paper all networks are rooted, so we will drop the word ‘rooted’.

This is a more general definition than sometimes used, in that it allows internal vertices to have any non-zero in-degree and any non-zero out-degree. A *phylogenetic tree* is a phylogenetic network in which all internal vertices have in-degree 1.

We draw a phylogenetic network with its single root at the top, its leaves at the bottom, and directed edges drawn with direction running down the page. Vertices of a phylogenetic network are called *tree vertices* if they have in-degree 1 (including leaves), and are called reticulate vertices (or reticulations) if they have in-degree greater than 1.

We say an internal vertex is *degenerate* if it either has in-degree and out-degree both equal to 1, or in-degree and out-degree both strictly greater than 1 (the former case are *degenerate tree vertices*, the latter are *degenerate reticulations*). A network is called *non-degenerate* if it contains no degenerate vertices.

A phylogenetic network is *binary* if the root has out-degree 2, and all internal vertices have total degree 3 (in-degree 2 and out-degree 1, or vice versa).

### 2.2. Trees, forests, and labellings

Phylogenetic trees and forests (collections of trees whose leaves partition the set {1, …, *n*}) have been shown to correspond to various partitions of finite sets. The basis for this correspondence is a labelling algorithm that, given a phylogenetic tree with leaves labelled 1, …, *n*, assigns labels to the internal vertices of the tree in increasing order from *n* +1 [6]. In the case of binary trees (those whose internal vertices have out-degree 2), there are a total of 2*n* – 2 labels, and a correspondence with perfect matchings follows by forming pairs of labels of sibling vertices (sibling vertices are those that share a parent) [4]. In the case of non-binary trees and phylogenetic forests, the same process of forming sets of sibling vertices gives a correspondence between the set of phylogenetic forests and all partitions of finite sets [10] (see Fig. 1).

**Figure 1.**
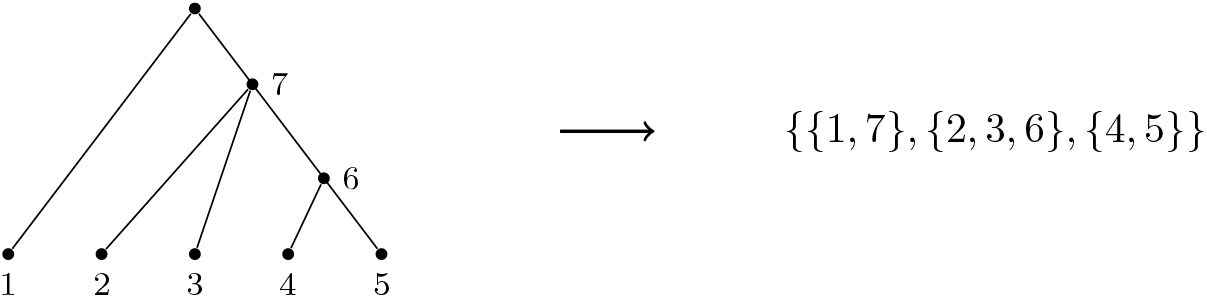
A tree that has been labelled according to the algorithm in [6], and its corresponding partition whose sets are sets of sibling vertices. This labelling algorithm is a special case of that given later for networks, in Algorithm 1.

In the next section we show how this labelling algorithm for trees and forests can be extended to some, but not all, phylogenetic networks. Those for which it can be extended we call ‘labellable’ networks.

## 3. Labellable networks

The labelling algorithm for binary trees given by [6] and [4], and extended to non-binary trees and forests in [10], takes a tree or forest with leaves labelled by [n], and progressively labels internal vertices in sequence, until all except the root are labelled. At each step the algorithm chooses a new vertex to label from those whose children are all labelled, by choosing the one whose children have the lowest label. Note, with trees and forests, the sets of children of vertices are disjoint.

In networks, sets of children of internal vertices are not, in general, disjoint, and it is necessary to define the following partial order to facilitate the labelling algorithm:

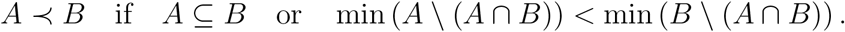

Note that this order reduces to the order used for labelling internal vertices in phylogenetic forests (in Algorithm 1 of [10]).

### Algorithm 1 Internal vertex labelling algorithm

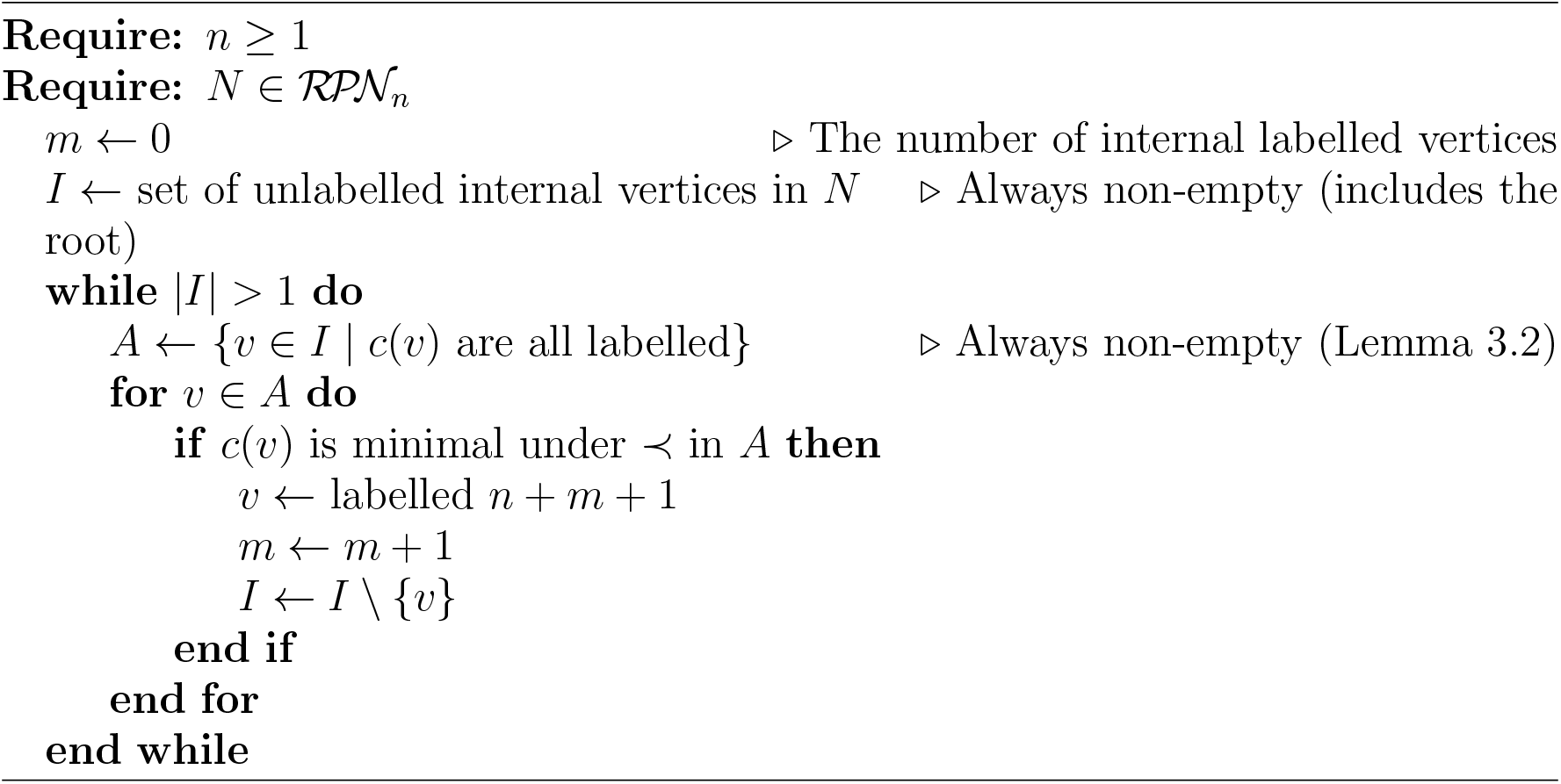

Algorithm 1 begins with a network whose leaves are labelled by [*n*] = {1, …, *n*} under the usual total ordering. At each step the algorithm takes a network that has been partially labelled, and chooses the minimal set of labelled sibling vertices (with respect to ≺) whose parent is not labelled, and assigns the next available integer to the parent. The output is a network whose (non-root) interior vertices are labelled by elements of *n* + 1, …, *n* + (#interior vertices) – 1.

Algorithm 1 is well-defined for a network *N* as long as a) it is always possible to find a set of sibling vertices whose parent is unlabelled, and b) each set of labelled siblings has only one parent. These are essentially existence and uniqueness style conditions.

### Definition 3.1.

A *labellable* network is one for which Algorithm 1 is well-defined.

While we show that a) holds for all phylogenetic networks (Lemma 3.2), it is easy for b) to fail (see Fig. 2).

**Figure 2.**
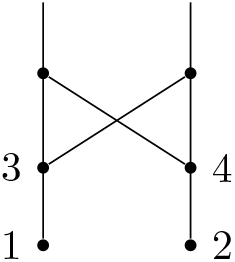
A network substructure that obstructs being labellable, since the vertices labelled 3 and 4 share the same set of parents.

### Lemma 3.2.

*Let N be a network with some vertices labelled, including all leaves. If N has any unlabelled vertices, then it contains at least one such vertex whose children are all labelled*.

*Proof*. For a vertex *v* in *N*, define *d*(*v*) to be the length of the longest path from *v* to a leaf. This is well-defined, since *N* is finite and acyclic.

For *v* an unlabelled vertex in *N*, if there is an unlabelled vertex *v*′ on a path from *v* to a leaf, then *d*(*v*′) < *d*(*v*). But *d* is bounded below by 1, for any unlabelled vertex (since the leaves themselves are already labelled). Therefore, repeating this process will eventually find an unlabelled vertex *u* for which all vertices on paths from *u* to a leaf are labelled. In particular, all of its children are labelled, as required.

Let *c_N_* (*x*) denote the set of children of *x* in *N*.

### Theorem 3.3.

*A network N is labellable if and only if c_N_*(*x*) ≠ *c_N_*(*y*) *for all vertices x* ≠ *y in N*.

*That is, N is labellable if and only if c_N_ is one-to-one*.

*Proof*. Lemma 3.2 shows that for any phylogenetic network the labelling algorithm will always find a set of labelled vertices whose parent is unlabelled. Let *U* be the set of such unlabelled vertices that are parents of labelled children. For the algorithm to be well-defined, it must be possible to use ≺ to place an order on the set *U*, via their sets of children.

The forward direction is immediate: if a network is labellable then it cannot have a pair of distinct vertices with the same set of children, since then those distinct vertices would not be able to be ordered.

Suppose for the reverse direction that sets of children of each vertex in *N* are distinct. Then any pair of unlabelled vertices with labelled children has distinct sets of children, and these can be ordered by ≺, thus allowing *N* to be labelled.

## 4. Covers and labellable networks

As with trees and forests, the labelling of non-root vertices in a network gives rise to a set of subsets of integers, namely the sets of sibling vertices (those sharing a common parent).

There are two differences between this set-up for networks and that in earlier work on forests. First, singleton sets may represent children of reticulations in networks, or indeed degree 2 internal vertices, instead of roots of trees. Second, the sets are not disjoint, because an integer labelling a reticulate vertex will have two sets of siblings. Consequently, sets of sibling vertices from a labellable network are not a partition of [m]. Instead, they are a ‘cover’ of [*m*], where a *cover* of [*m*] ≔ {1, ⋯, *m*} refers to a set of non-empty subsets of [*m*] whose union is [*m*]. Trivially, since a cover is a set, all of its member sets are distinct. The sets in a cover 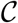 have a total order ≺ as defined above. We will denote the number of sets in 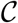 by 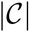.

We will call the set of sets of sibling vertices from a phylogenetic network *N the cover associated with N*. An example is given in Fig. 3. Note, repeat labels in a cover from a network correspond to reticulations.

**Figure 3.**
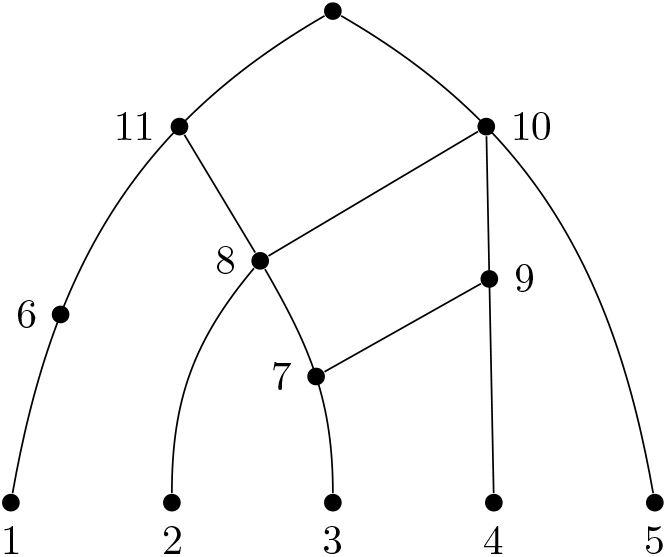
A labelled, degenerate phylogenetic network. This network has associated cover {{1}, {3},{2, 7},{4, 7},{5, 8, 9},{6, 8}, {10, 11}}.

Lemma 4.1 below is an analogue of a similar result for forests in the proof of [10, Theorem 6.5].

### Lemma 4.1.

*If* 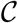 *is a cover of* [*m*] *associated with a phylogenetic network on *n* leaves, then*

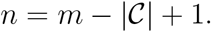

*Proof*. We can count the number of vertices in the network in two ways. If the number of sets in 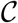 is 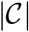, then there are 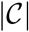 non-leaf vertices in *N*, and consequently 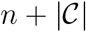 vertices overall. On the other hand, all vertices in the network are labelled by a unique element of [*m*], except the root. So there are [*m*] + 1 vertices in *N*. Thus, 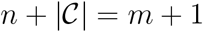, and so 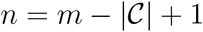 as required.

If a cover 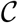 of [*m*] comes from a network, it also satisfies the properties in the following lemma:

### Lemma 4.2.

*If a cover* 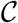 *comes from a labellable network N on *n* leaves, then*:

1. *The elements of* {1, ⋯, *n*} *are not repeated in* 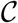; *and*
2. *For each* 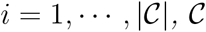 *contains at least i subsets of* [*n* + *i* – 1].

*Proof*. The leaves, labelled 1, …, *n*, have in-degree 1, and so each have precisely one parent, and are each members of precisely one set of siblings. Thus the labels of leaves are not repeated in 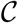, proving (1).

Claim (2) can be proved by induction on *i*. If *N* is labellable then in the first step there must be at least one set 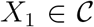 that is wholly contained in {1, …, *n*}. The label *n* + 1 is added to the vertex in *N* whose children are labelled by *X*_1_.

Suppose that for each *i* ≤ *k* there are at least *i* subsets of {1, …, *n* + *i* – 1} in 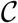. Then at each step of the labelling algorithm up to and including step *k*, a new vertex has been labelled whose children are labelled by the set *X_i_*, and in particular at step *k* the label *n* + *k* has been added to a previously unlabelled vertex in *N* whose children are labelled by *X_k_*.

Since *N* is labellable, after the *k*’th step there are *n* + *k* labelled vertices, and a set of labelled vertices *Y* whose parent is unlabelled. This set *Y* is distinct from *X*_1_, …, *X_k_*, since the sibling vertices labelled by those sets all have their parents labelled, by steps up to step *k*. Thus we have *X*_*k*+1_ ≔ *Y* is a subset of {1, …, *n* + *k* – 1, *n* + *k*} and we have *k* + 1 sets in 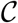 that are subsets of {1, …, *n* + (*k* + 1) – 1}, as required.

### Definition 4.3.

A cover satisfying the conditions in Lemma 4.2 is called an *expanding cover*.

### Theorem 4.4.

*The set of labellable phylogenetic networks is in bijection with the set of expanding covers*.

*Proof*. Lemma 4.2 shows that each labellable network gives an expanding cover.

For the reverse direction, if a cover is expanding, then a network can be deterministically constructed from the cover, using the following algorithm.

#### Algorithm 2 Network from an expanding cover

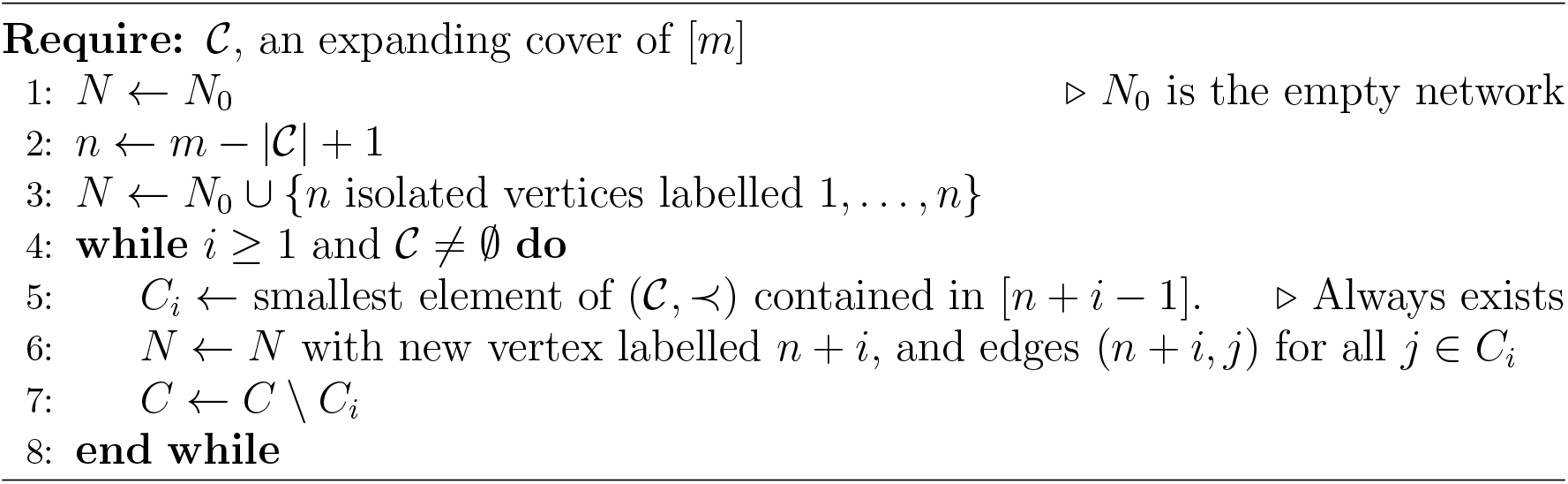

Note that in the intermediate stages the graph is not necessarily a phylogenetic network because it may not be connected, or may have several vertices of in-degree 0.

This algorithm is well-defined because of the properties of expanding covers — there will always be a set at the first step of the while loop (line 5) to choose — and it will terminate because 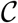 is finite and is reducing in cardinality by 1 with each iteration of the while loop at line 7. It will output a phylogenetic network because, with the exception of a single unlabelled vertex, every vertex has a label, and every labelled vertex is in a set in 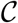 and so has a parent. The single unlabelled vertex is the unique root which has in-degree 0.

### Example 4.5.

Consider the cover:

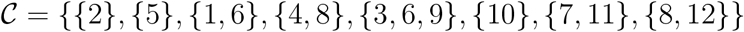

of [12], with 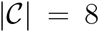. If this corresponds to a network, the network must have *n* =12 – 8 + 1 = 5, by Lemma 4.1.

It is expanding, because: it has two subsets of [*n*] = [5] (the definition of expanding requires at least one); three subsets of [6] (needs at least two); three subsets of [7] (needs three); four subsets of [8]; five subsets of [9]; six subsets of [10]; seven subsets of [11]; and eight subsets of [12].

See Fig. 4 for an illustration of the network constructed from this cover, using Algorithm 2.

**Figure 4.**
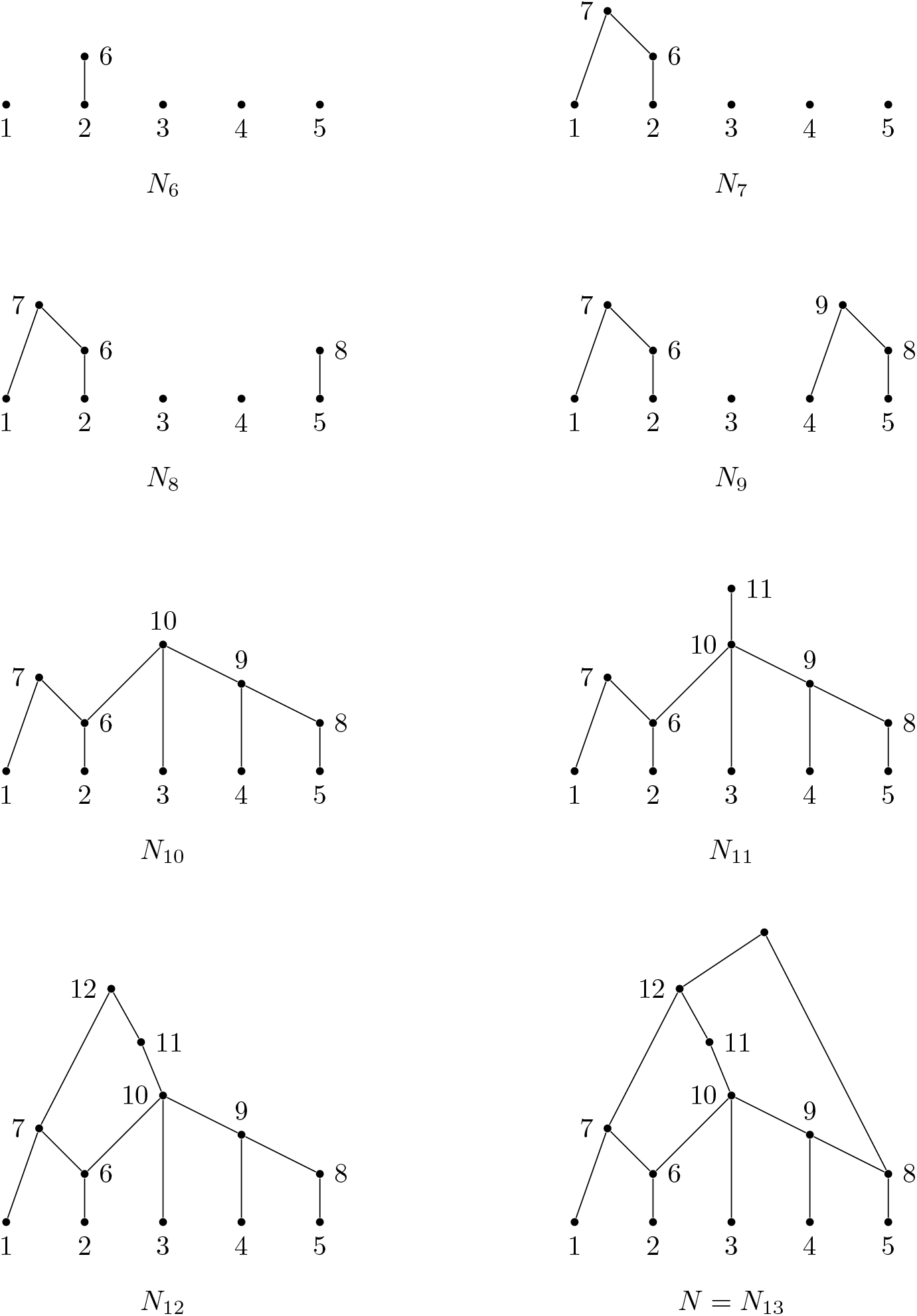
Construction of a network from the expanding cover 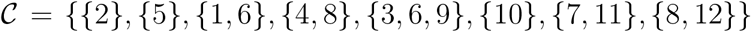 described in Example 4.5. The first step, consisting of 5 isolated vertices, is omitted. Note that the labels of the reticulate vertices (6 and 8) appear twice in 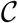.

The set giving rise to the vertex label *i* is determined by its position in an ordering of the sets in 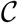. This ordering will use ≺ and is defined by the following algorithm:

1. For 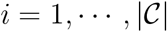, do:

a. *C_i_* is the minimal set in 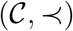 contained in [*n* + *i* – 1].
b. Set 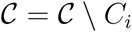.
2. Output sequence 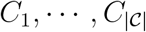.

Because this is the order that sets in a cover are used to assign new labels to vertices in the labelling algorithm (Alg. 1), we call it the ‘labelling order’ for a cover.

## 5. Covers and non-degenerate networks

The constraint on networks that they be non-degenerate requires an associated constraint on the corresponding set of covers. Non-degeneracy is equivalent to the condition that each internal vertex has either in-degree 1 or out-degree 1, but not both.

Let *N* be a non-degenerate labellable network with cover 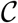. In terms of 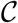, the in-degree of vertex *i* in 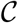 is precisely the number of sets in 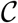 for which *i* is an element, while the out-degree of *i* is the size of the set in 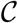 that gave rise to it. If *N* is non-degenerate, there must be corresponding constraints on the cover 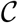.

With the sequence on the sets in 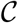 given by the labelling order, we now have the following two-part ‘order condition’ for each *i* ≥ 1:

- If |*C_i_*| > 1 then (*n* + *i*) appears *at most once* in 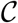; and
- If |*C_i_*| = 1 then (*n* + *i*) appears *more than once* in 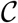.

### Theorem 5.1.

*The set of non-degenerate phylogenetic networks are in bijection with the set of expanding covers satisfying the order condition*.

*Proof*. For any phylogenetic network we have a unique expanding cover, by Theorem 4.4. If that network is non-degenerate, each vertex of out-degree greater than one must have in-degree equal to one, and each vertex of out-degree exactly one must have in-degree strictly greater than one.

If a vertex in a labellable non-degenerate network *N* has label *k*, then *k* = *n* + *i* for some *i* ≥ 1, since the leaves (labelled 1 to *n*) all have out-degree 0. This indicates that the label *k* was added by the *i*’th set *C_i_* in the labelling order on the expanding cover 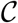 that corresponds to *N*.

If the set *C_i_* has cardinality one, then the vertex labelled *k* has out-degree one, since its children are the vertices whose labels are in *C_i_*. Since *N* is non-degenerate, the in-degree of the vertex labelled *k* must be strictly greater than one. This implies that *k* has more than one parent, and therefore more than one set of siblings. That is, *k* appears in more than one sets in 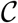, as required.

On the other hand if *C_i_* has cardinality strictly greater than one, then *k* has out-degree |*C_i_*| > 1, and since *N* is non-degenerate, it must have in-degree one. This is not possible if *k* appears more than once in 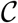, so it must appear just once, as required.

In the reverse direction, each expanding cover corresponds to a unique phylogenetic network by Theorem 4.4. If, in addition, the cover satisfies the order condition, then a) each vertex label that arises from a set *C_i_* of cardinality greater than one appears just once, and b) each vertex label that arises from a set *C_i_* of cardinality exactly one appears more than once. These constraints respectively force the vertex to be in-degree one and out-degree more than one, or in-degree more than one and out-degree one. In other words, the vertex is non-degenerate, and so the network as a whole is non-degenerate.

The order condition feels somewhat unsatisfying because it is not a passive property of the cover, but requires an additional order to be placed on it (the labelling order), and a check to be performed algorithmically. By contrast, the conditions for expanding covers are a static check for each 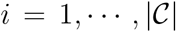. However, it is in fact ‘easy’ to check the order condition holds, for a given cover. The construction of the order on 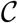 is linear in 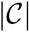, and the check of the order condition itself seems to be quadratic in 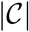 the check is done for each *i*, and for each check, membership is tested for each set in 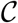. In other words, the whole process is at worst cubic in complexity.

### Example 5.2.

Recall the cover used in Ex. 4.5 was

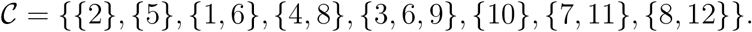

The labelling order on 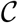 is as used in the networks construction in Fig. 4 (by design), namely ({2}, {1, 6}, {5}, {4, 8}, {3, 6, 9}, {10}, {7, 11}, {8, 12}). That is, *C*_1_ = {2}, *C*_2_ = {1, 6}, and so on, and with *n* = 5. To check the order condition here, we need to look at sets of size 1 and those of size > 1. The three of size 1 are *C*_1_ = {2}, *C*_3_ = {5}, and *C*_6_ = {10}. The order condition would therefore require 6 = *n* +1, 8 = *n* + 3, and 11 = *n* + 6 to appear more than once in 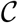. This is satisfied for 6 and 8, but not 11. This is seen in the resulting network by 11 labelling a degenerate vertex of in-degree and out-degree 1.

The sets of size > 1 are *C*_2_, *C*_4_, *C*_5_, *C*_7_, *C*_8_, and all of *n* + *i* for *i* = 2, 4, 5, 7, 8 appear at most once in 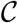, and so don’t give degenerate vertices. Note that the last of these, 13 = 5 + 8, does not appear at all, as the vertex added by that set is the last, which is the unlabelled root.

## 6. Labellable networks and other familiar classes

The property of being labellable defines a class of networks, and so it is natural to ask whether it is in fact one of the known classes, and if not, how it relates to the many well-studied classes of phylogenetic network. For overviews of the many known classes, we refer the reader to [16, Chapter 10] or [15].

Most of the well-studied classes are defined without permitting degenerate vertices (for instance orchard networks have been defined in the non-binary case [17], but not with degenerate vertices), so many results in this section are restricted to the non-degenerate case, whether binary or not.

Of the many classes previously defined, most are contained in the class of tree-based networks [11] — which are networks that have a spanning tree whose leaves are those of the network — so we first ask whether all labellable networks are tree-based.

It turns out that not all tree-based networks are labellable, because the property of being labellable excludes substructures such as that in Fig. 2, which are allowable in tree-based networks, as the following result makes clear.

### Corollary 6.1

(Corollary to Theorem 3.3). *If N is a nondegenerate tree-based network, then N is labelleable if and only if c_N_*(*x*) ≠ *C_N_*(*y*) *for all <u>tree</u> vertices x* ≠ *y in N*.

*Proof*. By Theorem 3.3, it suffices to show that for any non-degenerate tree-based network *N* the equality *c_N_*(*x*) = *c_N_*(*y*) cannot hold if one or both of *x, y* is reticulate. First suppose that *x* and *y* are both reticulate vertices. Then by the nondegenerate condition *x* and *y* each have exactly one (identical) child *z*, and again by the nondegenerate condition *z* is a reticulate vertex. However this stack of reticulations cannot exist in a tree-based network (by the antichain-to-leaf property described in [11]). Alternatively, suppose that *x* is a reticulate vertex and *y* is a tree vertex. Then (again by the non-degenerate condition) *x* has a single child, while *y* has at least two children and so *c_N_* (*x*) ≠ *c_N_* (*y*).

### Remark 6.2.

A further corollary of this last result is that if *N* is semiresolved (i.e. every tree vertex has outdegree 2) and tree-based, then *N* is labellable if and only if *N* is stable (this follows from Theorem 1 of [12], where the notion of ‘stable’ is defined).

In the case of *binary* tree-based networks, a tighter characterisation of those that are labellable is possible, as follows.

### Theorem 6.3.

*A binary tree-based network is labellable if and only if it has the property that p_N_* (*x*) ≠ *p_N_* (*y*) *for each pair of reticulate vertices x,y*.

*Proof*. Let *N* be a binary tree-based network.

Suppose that *p_N_*(*x*) = *p_N_*(*y*) for a pair of reticulate vertices. Since *N* is binary, *x,y* each have the same two parents (say *u,v*) and so (again since *N* is binary), *c_N_*(*u*) = *c_N_*(*v*). Hence, by Theorem 3.3, *N* is not labellable.

Conversely, suppose that *N* is not labellable. Then Theorem 3.3 implies that there exist vertices u,v with *c_N_*(*u*) = *c_N_*(*v*). Since *N* is binary this implies that either a) *c_N_*(*u*) = *c_N_*(*v*) = {*x,y*} for a pair of reticulate vertices *x,y*, and so *p_N_*(*x*) = *p_N_*(*y*), or alternatively b) *c_N_*(*u*) = *c_N_*(*v*) = {x}. But this second case can’t occur in a binary tree-based network, since it implies that *u, v*, *x* are three reticulations with arcs (*u, x*) and (*v, x*), which is impossible in a binary tree-based network. Thus *p_N_*(*x*) = *p_N_* (*y*) for a pair of reticulate vertices.

While not all tree-based networks are labellable, it is also true that not all labellable networks are tree-based. The substructure of a network shown at the right of Fig. 5 can be labelled, but is an obstacle to tree-based (it is a ‘zig-zag path’, as described in [20]).

**Figure 5.**
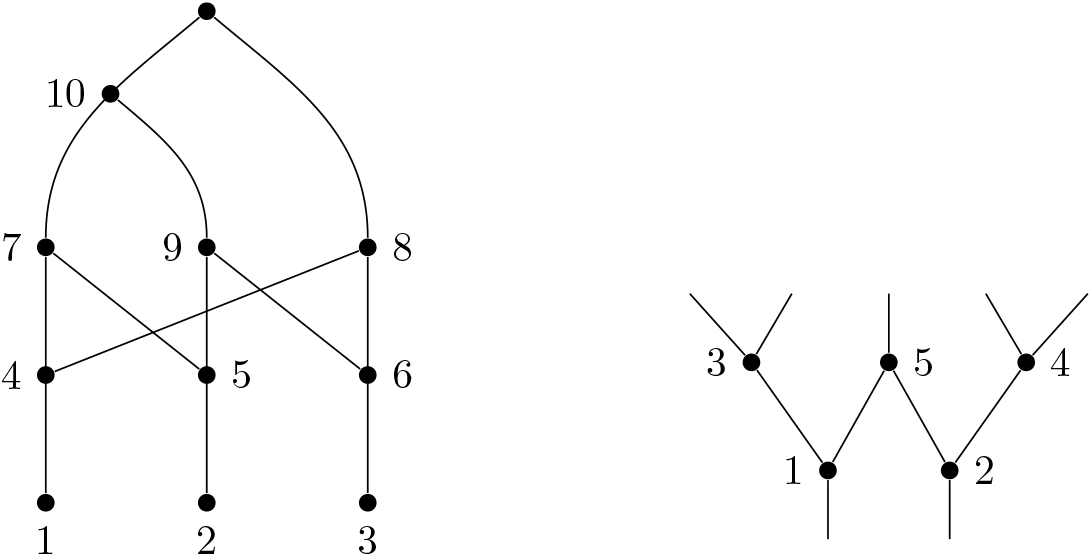
Left: A labellable network that is not orchard. It can be seen to be not orchard because it does not have any cherries or reticulated cherries. Right: A substructure called a zig-zag path [20] that prevents a network from being tree-based, but is nevertheless labellable.

**Figure 6.**
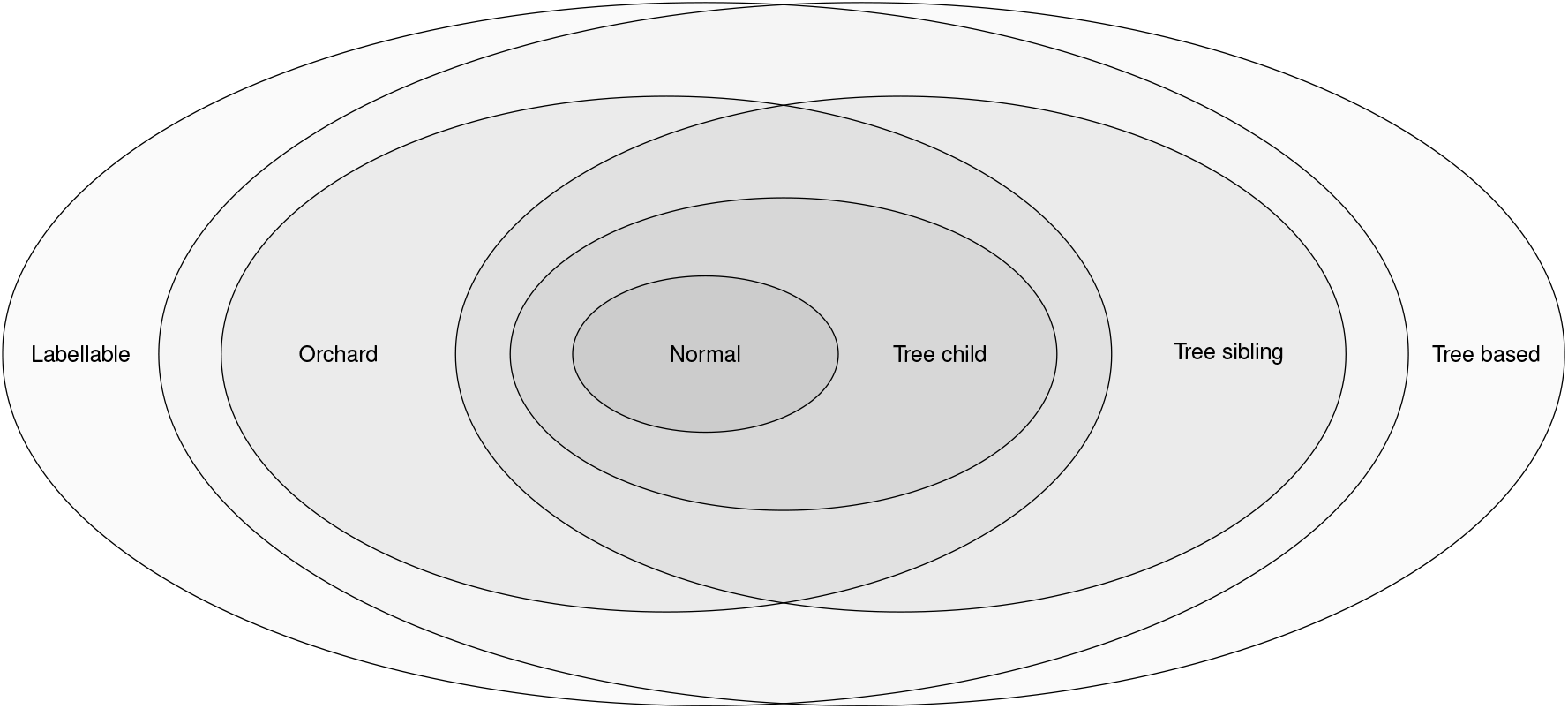
Relationships between labellable networks and some other classes. This figure applies for non-degenerate networks (non-binary permitted).

In summary, the classes of labellable and tree-based networks are not nested within each other.

In the remainder of this section we show that some other large classes that sit inside the tree-based networks — the orchard networks and tree-sibling networks — are also labellable. Thus, they sit in the intersection of the tree-based and labellable classes (see Fig. 6). Furthermore, other prominent classes of network are contained inside the intersection of orchard and tree-sibling. These include tree-child networks [14], and normal networks, which are tree-child (as also shown in Fig. 6).

*Orchard networks* are non-degenerate, rooted phylogenetic networks, that have the defining property that they can be reduced to a single point by a series of cherry and reticulated cherry reductions. These reductions replace a cherry or reticulated cherry with a simpler structure, reducing the network progressively. See [5] for the original definition and [17] for the extension to the non-binary case, and in particular a result we use in the proof below.

### Theorem 6.4.

*Orchard networks are labellable*.

*Proof*. Suppose that *N* is an orchard network, but is not labellable (we will derive a contradiction). Since every orchard network (binary or non-binary) is tree-based (by Corollary 4.5 of [18]), Corollary 6.1 implies that *N* has a pair of tree vertices u,v with *c_N_*(*u*) = *c_N_*(*v*) (and since *u,v* are tree vertices this shared set has size at least 2).

By Theorem 2 of [17] a non-degenerate network *N* is orchard if and only if some binary resolution of *N* (or *N* itself, if *N* is binary) admits an HGT-consistent labelling. However the vertices *u,v* with their shared set of children of size *k* ≥ 2 provides an obstruction to any resolution that allows a HGT-consistent labelling. Thus, *N* fails to be an orchard network, a contradiction.

### Corollary 6.5.

*If N is an orchard network then it does not contain any two vertices that have the same set of children*.

*Proof*. Since *N* is orchard it is labellable, by Theorem 6.4, and the result follows from the characterisation in Theorem 3.3.

Note, there are labellable networks that are not orchard, such as that in Fig. 5.

Theorem 6.4 also implies that tree-child networks [3] are labellable (since they are a subset of the orchard networks), but this can also be proved directly by looking at sets of children of vertices, as follows.

Recall that a vertex *v* in a network is *visible* if there is a leaf *i* ∈ [*n*] for which every path in *N* from the root to leaf *i* passes through v.

### Lemma 6.6.

*In any phylogenetic network, N, if c_N_*(*u*) = *c_N_*(*v*) *then neither u nor v are visible vertices*.

*Proof*. Suppose *c_N_* (*u*) = *c_N_* (*v*) and *u* is visible. Then there is a leaf *i* ∈ [*n*] for which every path *P* from *ρ* to *i* passes through *u*. Since *P* must also pass through one of the children of *u* (say *x*), let *P*′ be any path from *ρ* to *v*, and extends this path by adding edge (*v,x*) followed by the path used by *P* from *x* to *i*. This extended path is then a path from *ρ* to *i* that avoids *u*, contradicting the assumption that u is visible.

Note that a network *N* is tree-child if and only if every vertex is visible, and so an immediate consequence of Lemma 6.6 is that tree-child networks are labellable.

A phylogenetic network is *tree-sibling* if every vertex has a sibling that is a tree vertex [2]. In other words, every set of children (of a vertex in the network) has at least one tree vertex. Since a tree vertex is the child of exactly one parent, it is impossible for two sets of children to have the same parents, in a tree-sibling network. Thus, as an immediate corollary to Theorem 3.3, tree-sibling networks are also labellable.

### Corollary 6.7

(Corollary to Theorem 3.3). *Tree-sibling networks are labellable*.

Thus, the classes of orchard and tree-sibling networks are labellable, and other prominent classes the tree-child networks and normal networks [19] are both orchard and tree-sibling, and so sit in their intersection (and in particular, are also labellable). The above findings are summarized in Fig. 6.

## 7. Derived networks

In this section we return to the wider generality of allowing degenerate networks, and we show that every phylogenetic network has a quotient that is labellable.

Take any network *N* = (*V, A*) with leaf set [*n*] and define a relation ~ on *V* by *x* ~ *y* if and only if *c_N_* (*x*) = *c_N_*(*y*). Then ~ is an equivalence relation, and so *N* determines an associated network *N*′ = (*V*′, *A*′) where *V*′ are the equivalence classes of *V* under ~, and (*u, v*) ∈ *A*′ if and only if there exists *x* ∈ *u*, *y* ∈ *v*, with (*x, y*) ∈ *A*. Note that *N*′ has leaf set [*n*], and a single root.

In general, the network *N*′ may still have distinct vertices *u*, *v* for which 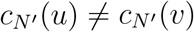, in which case *N*′ is not be labellable. Nevertheless, we can repeat the above process to construct a sequence N, *N*′, *N*″ ⋯ which stabilises after a finite number of steps at a network 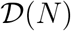 that has no distinct vertices *u*, *v* with 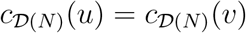, and thus 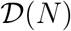 is labellable. Note that the number of steps required to reach this labellable network is finite since replacing an equivalence class (of vertices) of size at least one by a single vertex reduces the number of vertices in the network.

This provides a canonical (and idempotent) map from all networks to the class of labellable networks. An example of this map is given in Fig. 7.

**Figure 7.**
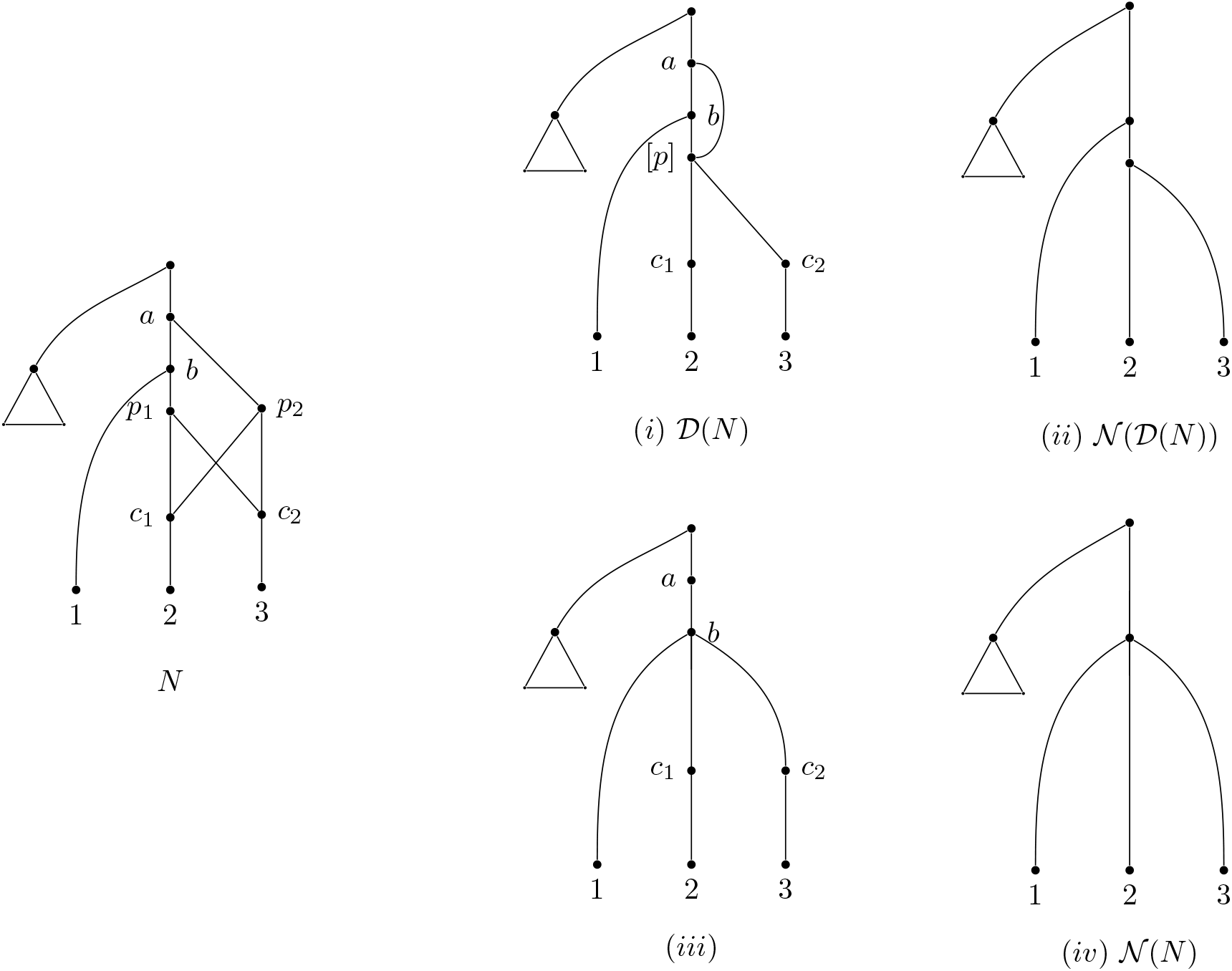
An unlabellable network *N* on the left, with the results of normalising, and taking the derived network. The pendant triangle represents an arbitrary tree. (*i*) shows its derived network 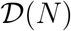, where [*p*] denotes the ~ equivalence class {*p*_1_, *p*_2_}, and (*ii*) shows the normalisation of that derived network, 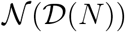. The bottom row shows the normalisation process described in [9]. (*iii*) shows the visible vertices and related edges (the first step of the normalisation process), and (*iv*) shows the complete normalised network 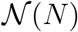 with degree 2 vertices suppressed. Note that 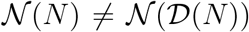 for this example.

### 7.1. Connection to normalisation

The above contains an echo of the normalisation map in [9], in that it is a map from the class of all phylogenetic networks to a specific class, in this case the labellable networks.

Recall we write 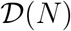 for the derived network of any network *N*, and let us write 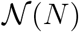 for its normalisation (as defined in [9]).

Note that normal networks are already labellable, so

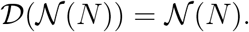

On the other hand, the normalisation of a network and of its labellable (derived) version need not be the same. That is, it is possible that

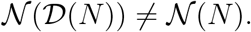

Thus we obtain two normalisations of the same network, one going via a labellization process. An example to illustrate this inequality is shown in Fig. 7.

## 8. Discussion

We have described a combinatorial correspondence for phylogenetic networks that maps to sets of covers of a finite set. Covers of finite sets that grow in a constrained way (the expanding covers) correspond to the large class of labellable phylogenetic networks. Those expanding covers that can be ordered in a particular way correspond to non-degenerate networks. The new class of labellable phylogenetic networks contains the class of orchard networks, but neither contains, nor is contained in, the class of tree-based networks. Because the class of labellable networks contains the orchard networks, it also contains the class of tree-child networks, and the class of normal networks.

There are a number of interesting further questions that remain to investigate.

For instance, we can describe when a *binary tree-based* network is labellable, based on the structure (Theorem 6.3). But are there conditions on the covers that force the network to be *orchard*, or *tree-child*, or *normal*?

It would also be interesting to more deeply understand the link between covers from degenerate and non-degenerate networks. A degenerate network can be made non-degenerate in a simple manner: we suppress any degree-two vertices, and blow up vertices of in-degree and out-degree more than 1 by replacing each such vertex by two vertices connected by an edge (so that a vertex of in-degree *d*_1_ and out-degree *d*_2_ is replaced by one vertex of in-degree *d*_1_ and out-degree one, connected to another of in-degree 1 and out-degree *d*_2_). What does removing the degeneracy mean for a cover, and what do these two degeneracy-removing actions involve at a cover level?

The connection between degeneracy-removal and covers extends to other actions on networks. Is it possible to describe actions such as nearest-neighbour-interchange (NNI) or subtree-prune-and-regraft (SPR) in terms of changes to the cover? And indeed this can be asked in the context of familiar families of network: can the cover of a network be manipulated to make it orchard, tree-child, or normal?

Finally, there may be connections to algebraic structures to explore. In the case of trees and forests, which correspond to partitions of finite sets, there is a corresponding set of partition diagrams that can be acted on by elements of the symmetric group or Brauer monoid [10]. What, if any, are the corresponding algebraic structures that correspond to covers, and can these be used to move around network space?

